# Optical coherence tomography of human fetal membrane sub-layers during dynamic loading

**DOI:** 10.1101/2023.03.09.531932

**Authors:** Kayvan Samimi, Emmanuel Contreras Guzman, May Wu, Lindsey Carlson, Helen Feltovich, Timothy J. Hall, Kristin M. Myers, Michelle L. Oyen, Melissa C. Skala

**Author notes:** **Correspondence:** Melissa C. Skala.

## Abstract

Fetal membranes have important mechanical and antimicrobial roles in maintaining pregnancy. However, the small thickness (<800 μm) of fetal membranes places them outside the resolution limits of most ultrasound and magnetic resonance systems. Optical imaging methods like optical coherence tomography (OCT) have the potential to fill this resolution gap. Here, OCT and machine learning methods were developed to characterize the *ex vivo* properties of human fetal membranes under dynamic loading. A saline inflation test was incorporated into an OCT system, and tests were performed on n=33 and n=32 human samples obtained from labored and C-section donors, respectively. Fetal membranes were collected in near-cervical and near-placental locations. Histology, endogenous two photon fluorescence microscopy, and second harmonic generation microscopy were used to identify sources of contrast in OCT images of fetal membranes. A convolutional neural network was trained to automatically segment fetal membrane sub-layers with high accuracy (Dice coefficients >0.8). Intact amniochorion bilayer and separated amnion and chorion were individually loaded, and the amnion layer was identified as the load-bearing layer within intact fetal membranes for both labored and C-section samples, consistent with prior work. Additionally, the rupture pressure and thickness of the amniochorion bilayer from the near-placental region were greater than those of the near-cervical region for labored samples. This location-dependent change in fetal membrane thickness was not attributable to the load-bearing amnion layer. Finally, the initial phase of the loading curve indicates that amniochorion bilayer from the near-cervical region is strain-hardened compared to the near-placental region in labored samples. Overall, these studies fill a gap in our understanding of the structural and mechanical properties of human fetal membranes at high resolution under dynamic loading events.

## 1 Introduction

The fetal membranes, also referred to as amniochorion or chorioamnionic membranes, are fetal tissues forming the amniotic sac composed of two closely adherent layers, amnion and chorion, consisting of several cell types, including epithelial, mesenchymal, and trophoblast cells embedded in a collagen matrix [1]. The amnion is the innermost layer and is in contact with the amniotic fluid and the fetus. The chorion, which is attached to the outer surface of the amnion, separates the amnion from the maternal decidua and uterus. A loose network of collagen and mucin forms the spongy layer that lies between the amnion and the chorion and allows them to slide against one another. Functionally, the fetal membranes support pregnancy by retaining amniotic fluid and protecting the fetus against infection. The fetal membranes normally rupture during labor. Premature (also referred to as prelabor) rupture of the membranes (PROM) is defined as rupture before the onset of labor and has a prevalence of nearly 10% in term deliveries [2]. PROM occurring before 37 weeks of gestation is referred to as preterm premature rupture of membranes (PPROM) and has a prevalence of nearly 3% of all pregnancies and 30% of PROM cases [2]. These conditions increase the risk of intrauterine and neonatal infection and associated complications, particularly when delivery is delayed following the rupture of membranes.

Prior studies of fetal membranes to establish mechanisms for structural alterations that lead to their rupture, either normally intrapartum at term or pathologically premature or preterm, point to the role of collagen matrix remodeling. Histological studies suggest that a “zone of altered morphology” (ZAM) forms in the fetal membranes overlying the cervix [3–5] characterized by swelling and disruption of fibrillar collagen network within the amnion (*i.e*., compact, fibroblast, and spongy layers) and thinning of cellular layers of the chorion (*i.e*., trophoblast and decidua). It has been hypothesized that the membranes in this zone have reduced tensile strength and become the initiating point of rupture. Two-photon and second harmonic generation (SHG) microscopy [6] has been used to investigate the structure of the fetal membranes and identify focal defects or “microfractures” in the sub-epithelial matrix that increase in size and density with increased oxidative stress and in PPROM cases [7]. The cause of PROM is likely multifactorial, involving an interplay of biophysical and biochemical pathways [8]. Molecularly, the degradation of collagen matrix is mediated by the balance of matrix metalloproteinase enzymes (MMP), hormones that affect their concentrations (*e.g*., progesterone and estradiol), and tissue inhibitors of metalloproteinase (TIMP), which normally shifts towards the end of gestation [9]. Intrauterine infection and host inflammatory response can alter this balance and increase the risk of PROM [10]. Mechanically, repeated stretching and uterine contractions can reduce the tensile strength of fetal membranes as evidenced by comparison of specimens from labored birth and unlabored cesarian delivery [11].

Studies of fetal membranes mechanics have investigated the correlation between membranes’ structural morphology and their break strength [12], changes in membranes strength with gestational age [13], and the difference between the mechanical properties of amnion and chorion [13]. Mechanical test methods employed include uniaxial tension [14–16], fracture and suture retention test [17,18], puncture test [19], planar biaxial tension [20], and biaxial inflation [11,21,22]. The inflation test geometry best represents the kind of loading that fetal membranes experience *in utero*. However, monitoring of tissue response to loading in these studies has been limited to contour mapping of the tissue surface from side view profiles acquired using an ordinary photography camera [21,23,24]. Thus, the response of individual sub-layers is not resolved. There is a need for a technique that can provide high-resolution and high-contrast imaging of fresh and unfixed fetal membranes under dynamic loading conditions and resolve the complex interactions of their constituting sub-layers. Such studies can inform computational biomechanical modeling of the fetal membranes and further our understanding of their modes of failure [25].

Optical coherence tomography (OCT) is an optical imaging technique akin to ultrasound imaging that uses low-coherence near-infrared or visible light to capture cross-sectional 2D or 3D images of the tissue with high (micrometer) resolution [26,27]. Previously, OCT was used to measure fetal membranes thickness [28,29] and identify features like atrophic chorionic ghost villi and chorionic pseudocysts [29,30]. However, these studies were limited to stationary samples without investigation of the effects of loading. Other studies combined speckle pattern interferometric thickness measurements with uniaxial loading to estimate the amnion rupture moduli in *ex vivo* samples [31]. Here, we combined inflation loading of fresh fetal membranes with cross-sectional OCT imaging to study their dynamic response to loading in a layer-resolved fashion at video frame rates. We identified collagen as a source of contrast in OCT images by comparison to histological sections and multiphoton microscopy, and trained machine learning semantic segmentation networks to identify amniochorion sub-layers in OCT images and estimate their thickness at every timepoint of the inflation loading experiments. Overall, these studies fill a gap in our understanding of the structural and mechanical properties of human fetal membranes at high resolution under dynamic loading events.

## 2 Materials and Methods

### 2.1 Sample acquisition and preparation

Fetal membranes and placentas were collected from normal term pregnancies between 37 and 41 weeks of gestation at the time of delivery. The study was deemed exempt human research by the institutional review boards of UnityPoint Heath Meriter Hospital and Intermountain Healthcare Utah Valley Hospital, and no active recruitment process was established. Two types of delivery were sampled. Placenta and membranes from labored vaginal deliveries (n=33) and unlabored elective Cesarian section deliveries (n=32) were collected at birth and placed in refrigerated phosphate-buffered saline (4°C PBS) before mechanical testing within 36 hrs.

Two anatomical regions were sampled for mechanical testing. A near-placental sample was obtained by cutting a 5-cm-diameter disk of membranes with an approximate 2 cm margin from the placental disk’s edge. Two near-cervical samples were obtained by cutting 5-cm-diameter disks of membranes close to either an ink-marked location (when marked by the surgeon during C-section delivery) or a best-guess cervical location based on presence of a clear fetal exit tear or maximum distance from the placental disk. One cervical sample was maintained as composite amniochorion (also referred to as chorioamnion in the literature), while the second sample was gently pulled apart to provide separate samples of the constituting amnion and chorion. The samples were maintained in room temperature PBS immediately before mechanical testing.

Smaller samples (1 cm squares) from the same near-cervical and near-placental locations were fixed in 4% paraformaldehyde (PFA) solution for multiphoton microscopy and histology preparation.

### 2.2 Mechanical testing via inflation under OCT imaging

Inflation tests were performed by laying the samples flat on a cylindrical 3D-printed resin loading vessel with an internal diameter of 30 mm and a flange width of 15 mm (Fig. 1). A clamping ring with the same dimensions and chamfered edges was placed over the sample and secured to the vessel using six equally spaced through-hole M4 bolts and nuts tightened to 3 Nm of torque. Once mounted, a 10-mm-wide uninterrupted margin of membranes on the flange supports the 30-mm-diameter open face of the sample disk. The resin surfaces were sanded rough to improve their grip. A constant-flow syringe pump connected to an open water column at the base of the vessel provided a nearly linear saline pressure ramp at a rate of 10 kPa per minute until rupture. The OCT imaging scan head and objective lens were positioned co-axially above the mounted sample disk and transversely imaged the diameter of the inflating sample at a rate of three frames per second. Whenever the sample apex reached the top of the OCT imaging depth range (2.65 mm), the pressure ramp was paused and the OCT scan head was moved up to allow additional inflation range, and the pressure ramp was resumed within a few seconds. An inline pressure sensor measured the applied saline pressure at every time point. The inflation test was performed for the composite amniochorion samples from the near-placental and near-cervical regions, as well as the individual amnion and chorion samples from the near-cervical region.

**Fig. 1.**
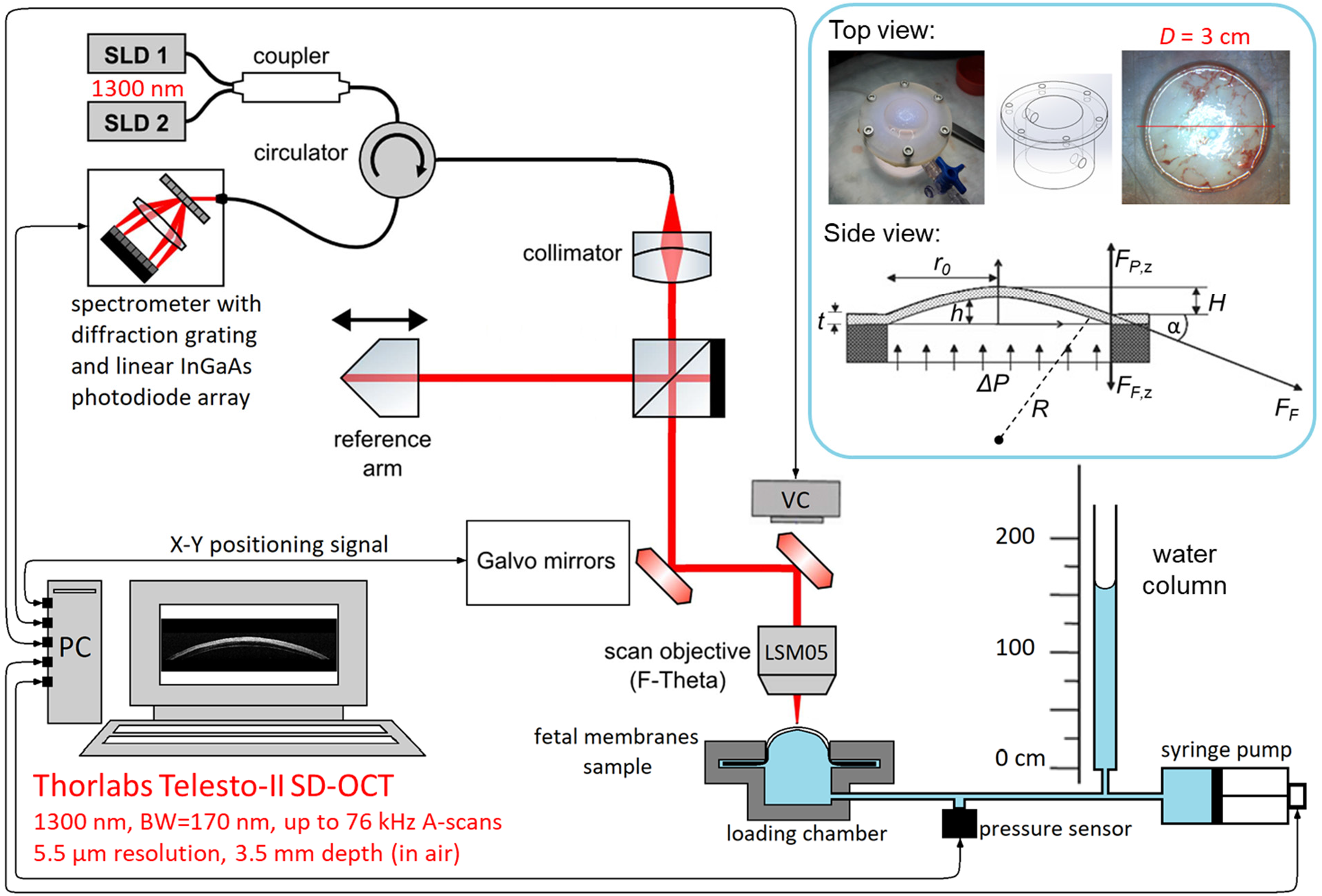
Fetal membranes mechanical testing via inflation with normal saline. Disks of fetal membranes (diameter = 50 mm) were mounted onto a cylindrical loading chamber (internal diameter = 30 mm, flange width = 15 mm) and secured with a clamping ring, resulting in an inflatable exposed disk of membranes with a 30 mm diameter. A constant flow syringe pump connected to an open water column at the base of the loading chamber provided a nearly linear saline pressure ramp at a rate of 10 kPa per minute until rupture. The OCT imaging scan head was positioned co-axially above the sample and transversely imaged the diameter of the sample disk at a rate of 3 frames per second. An inline pressure sensor measured the applied pressure at every timepoint. Inset shows the top view schematic and photograph of the loading chamber as well as the side view schematic and force diagram of the sample in the loading chamber. *ΔP* is the applied saline pressure. *H* is the sample’s apex rise. *r_0_* is the sample disk radius. *R* is the radius of curvature for the inflating sample. *t* is the sample thickness. *F_F_* is the frame force applied to the sample at the clamping edge. *α* is the angle between the sample and the flange. *F_P,z_* is the vertical component of the fluid force on the sample at the clamping edge. *F_F,z_* is the vertical component of the frame force on the sample at the clamping edge. SLD, superluminescent diode; VC, video camera; BW, bandwidth; SD-OCT, spectral domain optical coherence tomography.

OCT imaging was performed using a commercial spectral-domain system (Telesto-II, Thorlabs) equipped with a super-luminescent diode pair light source centered at 1300 nm with a bandwidth of 170 nm and a spectrometer with diffraction grating and a 2048-pixel linear photodiode array camera. Corresponding axial resolution and imaging depth in water were 5.5 μm and 2.65 mm, respectively. An LSM05 telecentric scan lens (Thorlabs) with effective focal length of 110 mm and working distance of 94 mm was used, providing a large field of view of 31 mm diameter and a beam spot size of 33 μm. Galvo mirror scan step size was set to 10 μm. The acquired spectral data were post-processed (including K-space linearization, dispersion compensation, and apodization with a Hann window function) before being Fourier transformed to produce intensity images with a pixel size of 10 μm both axially and laterally (3100 × 265 pixels in B-scans). The resulting images were saved as multipage TIFF files.

### 2.3 Multiphoton microscopy of fixed samples

Unstained fixed samples were imaged on a custom multiphoton microscope built around an Eclipse Ti-E inverted microscope (Nikon Instruments) using a titanium:sapphire tunable femtosecond-pulsed laser source (InSight DS+, Spectra Physics) and a 20×/1.0NA WI objective lens (Zeiss). Emissions were filtered using bandpass filters (Semrock) 550/100 nm for channel 1 and 440/80 nm for channel 2 before detection by photomultiplier tubes (H7422P-40, Hamamatsu). The excitation laser was tuned to a wavelength of 905 nm with an average power between 5 and 10 mW at the sample. Two-photon excited cellular autofluorescence was collected in channel 1 and second harmonic generation of collagen was collected in channel 2. Image field of view was 500 μm × 500 μm and the image pixel count was 512 × 512. A z-stack with steps of 1 μm was collected using the motorized sample stage spanning the thickness of the amniochorion sample. The z-stack volume was resliced in post-processing to produce cross-sectional images of the amniochorion layers that are comparable to histology cross-sectional preparations.

### 2.4 Histology

Fixed amniochorion samples were paraffin embedded and three consecutive microtome cross sections with 5 μm thickness were stained with standard hematoxylin and eosin (H&E), picro-sirius red for collagen, and Masson’s trichrome for contrast between nuclei, cytoplasm, and collagen. Stained sections were imaged using a brightfield microscope (Aperio ImageScope, Leica) with 20× magnification.

### 2.5 Convolutional neural network image segmentation

To enable assessment of changes in sub-layer thicknesses during loading based on video-rate OCT images (consisting of hundreds of B-scan frames per loading test), an automated semantic segmentation algorithm was implemented that identifies constituting layers of amniochorion. A fully convolutional neural network (CNN), ReLayNet [32], that was originally proposed for segmenting retinal layers and edema in ophthalmic OCT images was retrained using randomly selected and hand-segmented B-scan frames (n=242) of composite amniochorion samples under various inflation loads. Images were manually segmented in ImageJ by experienced observers using OCT image landmarks identified through comparison to histology and microscopy images of amniochorion. Image pixels were assigned one of six distinct labels: Four tissue layers for fetal amnion, spongy layer, chorion, and maternal decidua, and two non-tissue layers for saline below and air above the sample. The hand-segmented dataset was expanded (by a factor of five) using standard geometric image transformations such as scaling, cropping, rotation, translation, and horizontal mirroring, to create an augmented dataset. Ten-fold cross-validation was performed by splitting the augmented hand-segmented dataset into ten subsets. The CNN was trained ten times, leaving one subset out as the test dataset each time. Segmentation performance was evaluated using the Dice coefficient of these test images, defined as two times the area of the intersection of ground truth mask and CNN-generated mask, divided by the sum of the areas of both masks. Then the CNN was used to segment OCT images of complete inflation experiments and create corresponding layer masks.

Mean thickness of each layer for every frame was calculated in post processing by dividing the area of the layer mask by its length. The layer length was calculated as the length of a second-degree polynomial curve (i.e., parabola) fitted to the layer mask.

### 2.6 Analysis of inflation loading test data

Upon application of saline pressure, the membrane sample bulges into a pseudo-spherical shape. Given the saline pressure readouts from the sensor and cross-sectional OCT images of the sample at every timepoint, the loading curve for each inflation test is attainable. First, sample deformation was estimated from the OCT images by tracking the bulging sample’s apex rise between consecutive OCT frames using a custom image registration code in MATLAB (MathWorks) maximizing the 2D cross-correlation of a region of interest (ROI) containing the sample apex between frames. Knowing the fixed geometry of the loading chamber and clamping ring (Fig. 1), this measured dome apex rise can be converted to a measure of strain. We estimate strain as

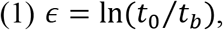

where *t*_0_ is the initial thickness of the membrane and *t_b_* is the thickness of the bulged membrane. For simplicity, we assume that the bulged membrane is spherical with a radius of curvature, *R*, and that the material is incompressible. As such, the total volume of the bulged membrane is assumed constant across timepoints. The radius of curvature can be calculated from the apex rise, *h*, and the radius of the sample disk, *r*_0_ (which is 15 mm in our experiments), as 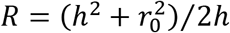. Equating the initial and the bulged membrane volumes, 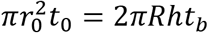, allows us to estimate the strain from (1) as:

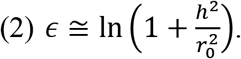

Given the slow inflation of the sample, we can estimate the membrane tension and stress from the equilibrium of counter-acting vertical forces on the membrane, from the pressurized saline on one side, and from the frame forces at the clamping ring edge, at any timepoint. This equilibrium can be written as 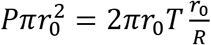, where *P* is the saline pressure, *T* is the membrane tension, and *r*_0_/*R* gives the sine of the angle that the membrane makes with the clamping edge of the frame. From this equation, the membrane tension and stress can be estimated as:

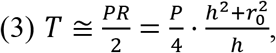

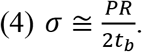

However, (4) assumes that the composite amniochorion is uniformly stressed along its thickness, which, given the non-uniform composition of collagen matrix across amniochorion layers, is likely not a valid assumption. Therefore, in all analysis in this work, we refrain from converting the tension estimates to stress estimates, and analyze tension-strain loading curves instead of stress-strain curves. Accordingly, we only report tangent stiffness in [N/m] instead of Young’s moduli in [Pa].

### 2.7 Statistical analysis

Statistical analysis and graph creation were done in MATLAB (MathWorks). Pearson’s correlation coefficient was used to evaluate presence of linear correlation, or lack thereof, between morphological measurements of the samples and their loading test outcomes.

Significance of differences in morphological or mechanical measurements between different sample groups (*i.e*., near-cervical or near-placental sampling locations) was determined using the two-tailed paired sample *t*-test.

## 3 Results

### 3.1 OCT intensity imaging is sensitive to collagen gradients in amniochorion layers

Prior studies have defined the layers of the amniochorion with histology [28–30] and two-photon endogenous microscopy (2P microscopy) [6,7]. Therefore, histology and 2P microscopy were performed in paired samples with OCT volume imaging under mechanical loading to identify sources of contrast in the OCT volumes. Cross-sectional views of 2P microscopy z-stacks (Fig. 2) show cellular autofluorescence (in red) and second harmonic generation of collagen (in green) which characterize the constituting layers of the amniochorion. From the bottom, amnion consists of a monolayer of epithelial cells followed by the highest density of collagen in its basement membrane and compact layer (compared to any other layer). Next, the amnion’s fibroblast cell layer is seen within its collagen matrix. A mostly acellular loose network of collagen and mucin forms the spongy layer that separates the amnion from the chorion and allows them to slide against one another. The spongy layer can accumulate fluids and shows large variability in thickness. The reticular layer of the chorion presents with chorionic mesenchymal cells in the collagen network ending with a more densely collagenated pseudo-basement membrane at the interface with the trophoblast layer. The trophoblast layer is the outermost layer of chorion that consists of several layers of trophoblast cells that are in contact with maternal decidua. This layer does not present with collagen, in contrast to the reticular layer below and maternal decidua above. Histological staining for collagen confirms the observed gradients in collagen content across amniochorion layers in two-photon microscopy images, with the amnion showing the highest density of collagen, the spongy layer showing a loose network of collagen fibers, and the trophoblast layer showing a distinct collagen-free band of cytotrophoblasts with an epithelial-like structure, followed by maternal parietal decidua cells and collagen (which the trophoblasts invade).

**Fig. 2.**
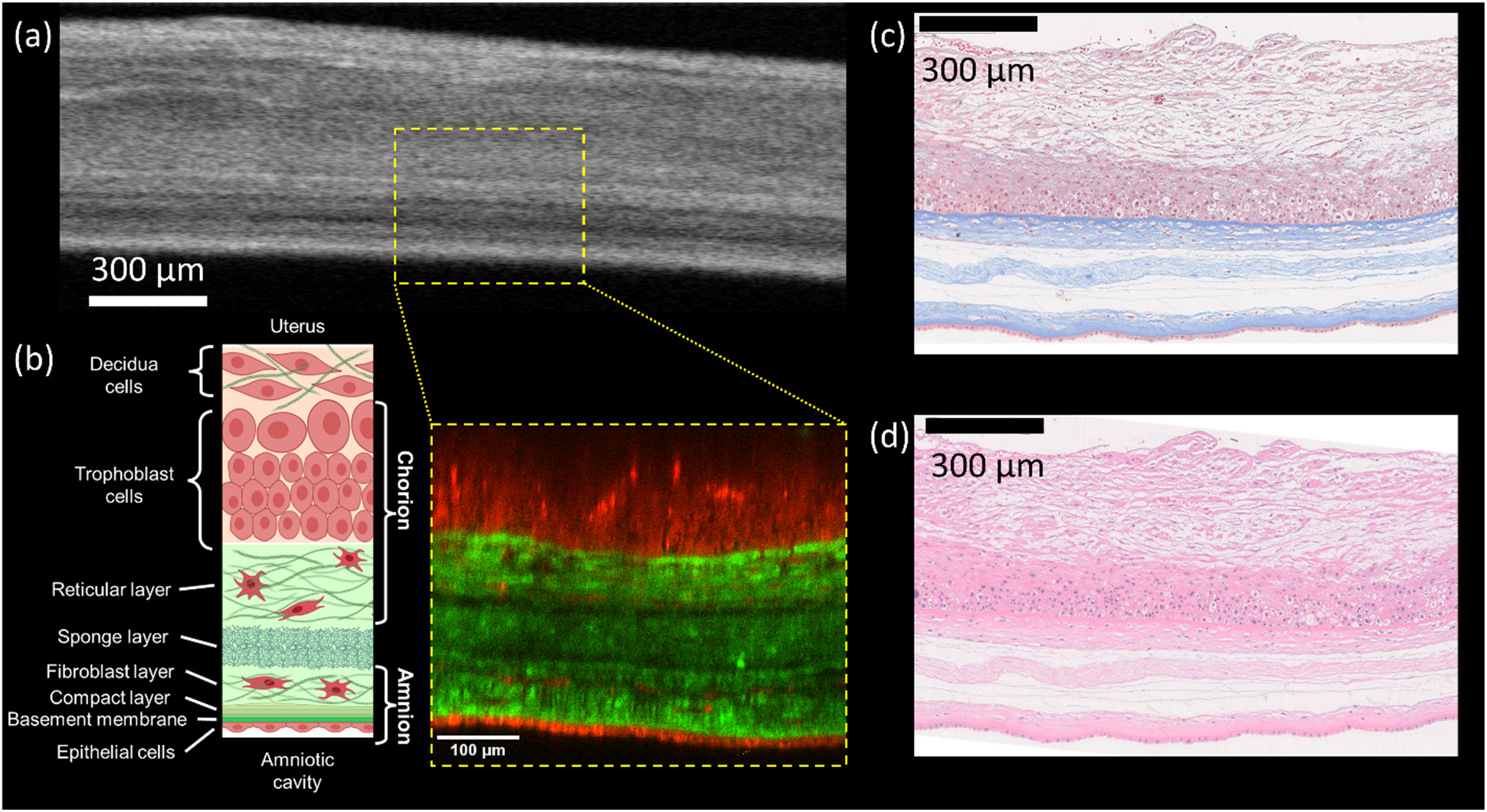
The multilayered amniochorion presents with cellular and acellular layers with gradients in collagen content which produce contrast in OCT images. Samples of human fetal membranes acquired from a term C-section delivery at 39 weeks and 2 days of gestation were imaged using optical coherence tomography, two-photon fluorescence microscopy, and bright field microscopy after histological staining. (a) OCT image visualizes the constituting layers of the amniochorion, with dense collagen layers and collagen fibers generating higher backscattering and appearing brighter while water and air gaps appear darker. (b) Illustration diagram and cross-sectional view of a two photon-excited autofluorescence microscopy Z stack of fetal membranes acquired in two spectral channels. Femtosecond-pulsed excitation laser was tuned to 905 nm and cellular autofluorescence was captured using a 550/100 nm emission filter (shown in red) and second harmonic generation of collagen was captured using a 460/50 nm emission filter (shown in green). Amnion (bottom) presents with an epithelial monolayer followed by a compact collagen layer and fibroblast layer. A loose network of collagen forms the spongy layer that separates and allows amnion to slide against chorion. Chorion consists of a collagen-rich reticular layer with sparse stromal cells followed by collagen-free and densely packed trophoblast cells. Decidua are maternal cells from the uterine lining that are fused to the outer fetal trophoblast layer. (c) Histology section of the amniochorion with Masson’s trichrome staining shows presence of collagen (in blue) in the basement and mesenchymal cell layers of amnion and chorion as well as the spongy layer in between. Cells cytoplasm are stained pink and cells nuclei are stained brown. The trophoblast layer of the chorion presents a collagen-free band of densely packed trophoblast cells that is distinct from the collagenated layers of chorion basement below and decidua above it. (d) H&E staining shows the nuclei and cytoplasm in the cellular layers of amnion and chorion. Comparison with two-photon microscopy and histology images suggests collagen is a source of contrast in OCT images, and that the constituting layers (i.e., amnion, spongy, chorion, decidua) can be differentiated on OCT intensity images based on the identified landmarks.

Comparison of OCT intensity images with corresponding 2P microscopy and histology images (Fig. 2) reveals collagen density as a source of contrast in OCT images. Dense collagen layers and collagen fibers generate high backscattering and appear bright, while water and air gaps are less backscattering and appear dark on OCT intensity images. The collagen-free band of cytotrophoblast cells in the chorion produces a low-intensity band on OCT images that stands in contrast to its collagen-rich basement membrane in the reticular layer below which has higher OCT intensity. The spongy layer with its loose collagen network and high water content typically presents with lower intensity on OCT images. However, due to its compressibility and perfusion, its OCT intensity tends to increase as its thickness decreases under load. The amnion generates high OCT signal intensity and stands out against the lower-intensity spongy layer above and the zero-intensity saline below it. This characterization of OCT intensity images of composite amniochorion allows for segmentation of individual layers on video-rate OCT images acquired during inflation loading tests and observation of associated dynamics.

### 3.2 Automated CNN segmentation of OCT images reveals dynamic morphological changes during inflation loading

The OCT image landmarks identified through comparison to 2P microscopy and histology were used to inform manual segmentation of a randomly selected OCT image set of amniochorion samples under various inflation loads between 0 and 20 kPa (n=242 images) for training and testing of the semantic segmentation CNN. Ten-fold cross-validation was performed by splitting the hand-segmented dataset into ten subsets. The CNN was trained ten times, leaving one subset out as the test dataset each time. The Dice coefficient metric was calculated between the CNN-generated masks and human-segmented masks in the test set. The mean and standard deviation of the achieved Dice coefficients for the four amniochorion tissue layers were 0.83 ± 0.04 for the amnion, 0.84 ± 0.11 for the spongy layer, 0.78 ± 0.10 for the chorion, and 0.91 ± 0.05 for the decidua. Fig. 3 shows a representative amniochorion OCT intensity image along with its corresponding CNN-generated layer mask overlays, along with the test image set mean Dice coefficients for each layer.

**Fig. 3.**
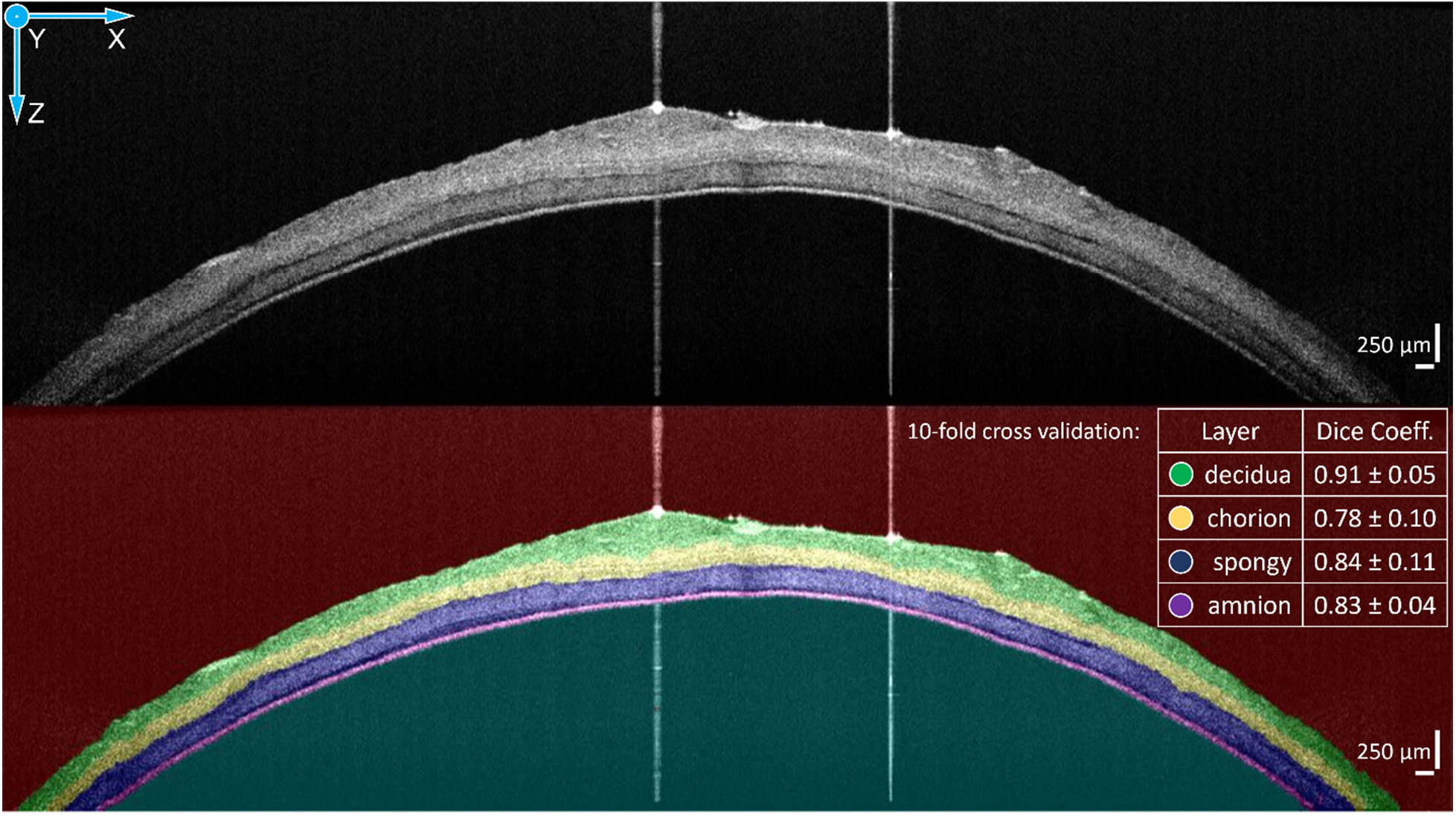
Convolutional Neural Network can segment amniochorion layers on OCT images in good agreement with human expert. A fully convolutional neural network from retinal OCT literature (ReLayNet [32]) was retrained using hand-segmented OCT images (n=242) of fetal membranes acquired during inflation loading tests. 10-fold cross validation was performed by splitting the hand-segmented images into ten subsets. The CNN was trained ten times by leaving one subset out as the test dataset. The agreement of the CNN segmentation output with the human segmentation was measured using the Dice coefficient metric on the test dataset. The automated CNN segmentation performs well with Dice coefficients of 0.8 or better for all constituting layers of the amniochorion.

The trained CNN was used to segment entire inflation loading test OCT image sets and the CNN-generated masks were post-processed in MATLAB to extract mean layer thickness values for each of the four layers at every timepoint, as described in the Methods section. Additionally, sample apex rise was estimated from the images and used, along with the sensor pressure readings, to generate loading curves. These values were then converted to strain and tension estimates to generate the tension-strain loading curve and calculate the slopes (*i.e*, tangent stiffness) at low- and high-load portions of the inflation test. Fig. 4 shows a representative inflation loading experiment and its associated loading curves and layer thicknesses over time. Supplemental videos (Visualization 1-2) show the complete inflation loading time course for near-cervical and near-placental samples, respectively, from the same delivery (labored natural birth at 39 weeks and 1 day of gestation).

**Fig. 4.**
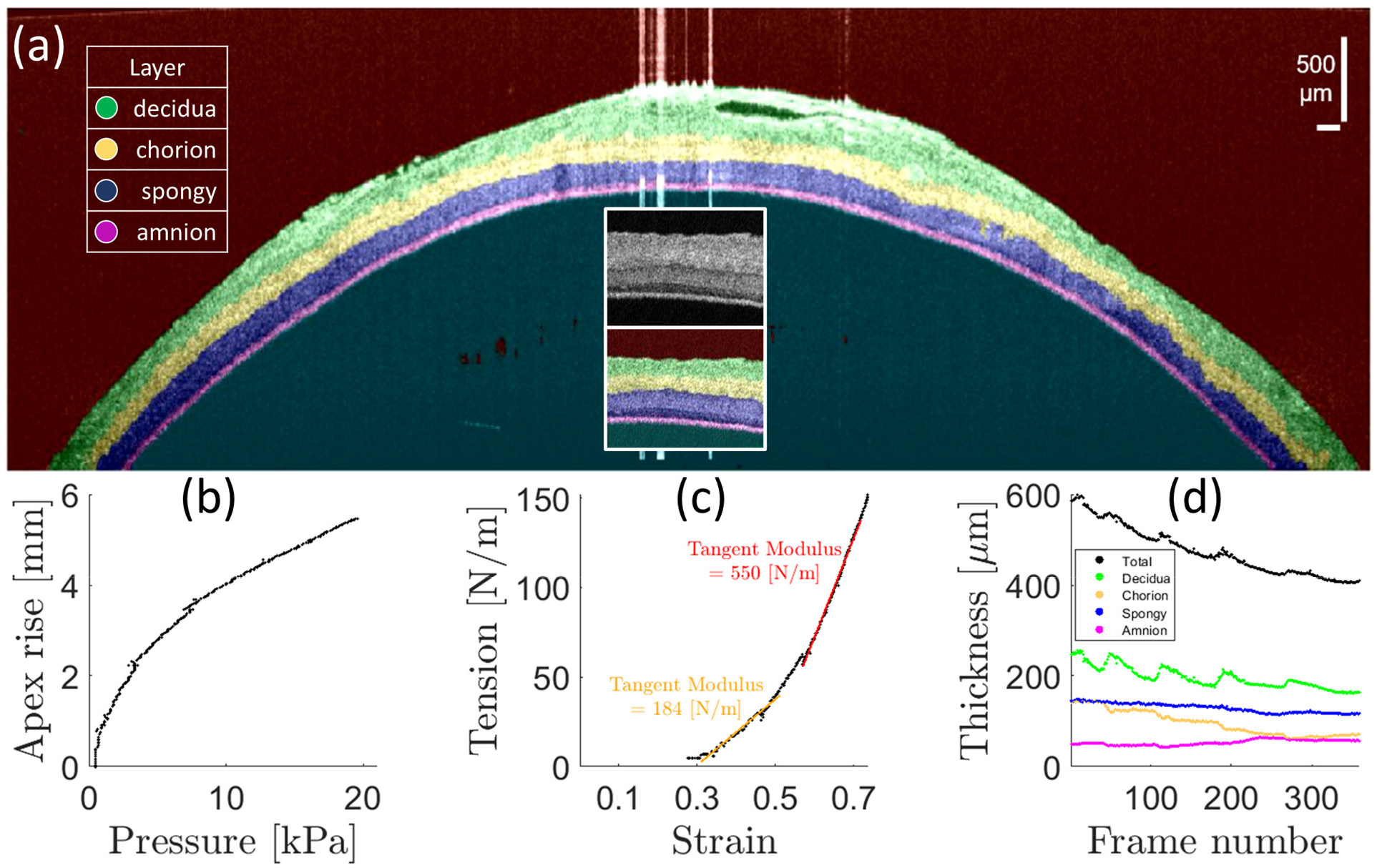
Loading curve and layer thicknesses can be extracted from OCT images of the inflation test. (a) Representative segmented OCT image frame from an inflation loading test. Inset shows raw intensity and layer-segmented images using a convolutional neural network. (b) Tissue deflection (apex rise) plotted against applied saline pressure at every time point. (c) Sample loading curve (tension vs. strain) derived from apex rise and pressure measurements. (d) Average thickness of individual layers plotted over time (see Visualization 2 for the complete inflation time course).

### 3.3 Layer-specific OCT inflation test confirms that the amnion is the major load-bearing layer in fetal membranes

Next, we confirm that the layer-specific OCT inflation test is consistent with prior work [13] that showed that the amnion is the major load-bearing layer in fetal membranes. The inflation pressure at the moment of rupture is plotted for composite amniochorion, separated amnion, and separated chorion samples from the near-cervical region in both the labored delivery and unlabored C-section birth groups (Fig. 5(a)). Using paired *t*-tests, we found that the rupture pressure for amnion alone is not significantly different from the composite amniochorion, while the chorion alone ruptures at a significantly lower pressure (nearly half) compared to the amnion or composite amniochorion. Therefore, while amnion is the thinnest constituting layer of the fetal membranes (<100 μm), it is also the major load-bearing layer responsible for the bulk of the amniochorion’s strength.

**Fig. 5.**
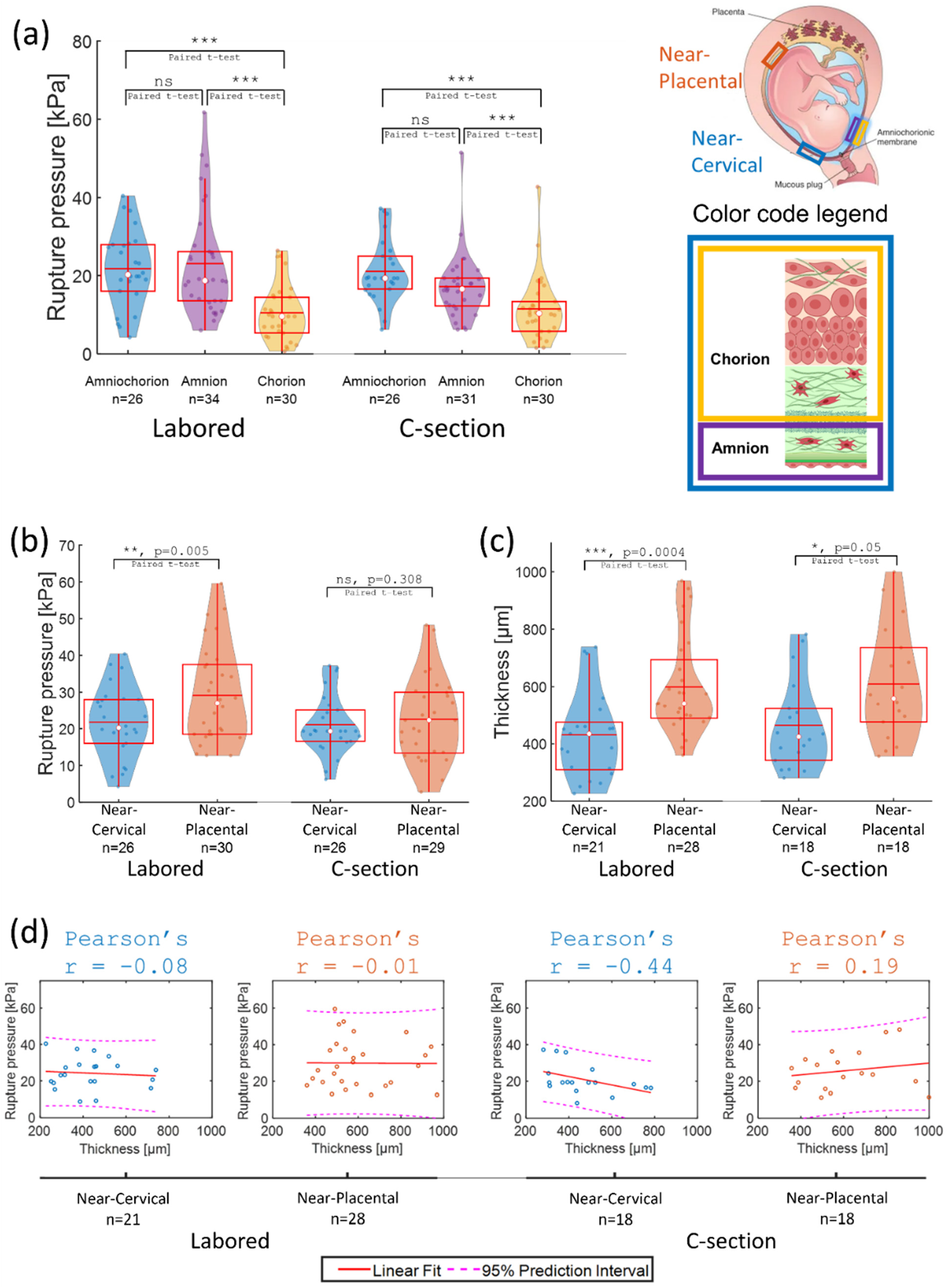
(a) Amnion is the major load-bearing layer. For both the labored and the C-section delivery groups, intact amniochorion bilayer and separated amnion and chorion were sampled from the near-cervical region and subjected to saline inflation until rupture. Paired *t*-test shows that the rupture pressure of amnion alone is not significantly different from the composite amniochorion. However, the chorion ruptures at significantly lower pressures than either amnion or amniochorion. Despite being thinner, amnion is the major load-bearing layer of the fetal membranes. **(b,c) Strength and thickness are location dependent.** Composite amniochorion was sampled from the near-cervical region and the near-placental region in both labored and C-section delivery groups. The two samples from each delivery were subjected to the inflation test and their rupture pressure and total thickness were compared against one another. (b) The rupture pressure of the near-placental sample was significantly higher than the near-cervical sample in the labored delivery group. However, the difference did not reach statistical significance in the C-section delivery group. The rupture pressure of the near-placental samples in the C-section group was also lower than the labored group. (c) The total thickness of the amniochorion in the near-cervical region is significantly lower than the near-placental region for both labored and C-section delivery groups. **(d) Thickness does not predict strength.** Total thickness of the amniochorion does not correlate strongly with the rupture pressure of the fetal membranes in any of the sampled regions or delivery groups. This finding is consistent with the results of individual amnion and chorion inflation loading tests that suggest amnion is the major load-bearing layer and that other layers, despite their higher and variable thickness, contribute minimally to the load-bearing strength of the composite amniochorion. These results are consistent with prior reports [12]. Box plots show the interquartile range and whiskers extend from the box to 1.5 times the interquartile range. White dots show the median and red horizontal lines show the mean.

### 3.4 Strength and total thickness of fetal membranes are reduced near the cervical os, but the two measures are not correlated

Location dependence of morphology and mechanical strength of the composite amniochorion was investigated by comparing samples from the near-cervical region against samples from the near-placental region in both the labored delivery and the unlabored C-section groups. Fig. 5 (b) shows the rupture pressure plots. The near-cervical samples have lower rupture pressure than the near-placental samples in the labored delivery group (*p* = 0.005, paired *t*-test). However, the difference does not reach statistical significance in the C-section group (*p* = 0.308). Fig. 5 (c) shows the total thickness of the amniochorion samples. The near-cervical samples are significantly thinner than the near-placental samples in both labored (*p* = 0.0004) and C-section (*p* = 0.05) delivery groups.

The correlation between total thickness and amniochorion strength was investigated in Fig. 5 (d) by performing a linear regression analysis. We found that a decreased total thickness of the amniochorion is not coincident with a reduced rupture pressure in either of the sampling regions or birth groups and that, consequently, total thickness is not a predictor of fetal membranes strength. Fig. 6 takes a closer look at the breakdown of constituting layer thicknesses between near-cervical and near-placental samples as measured from the CNN-segmented OCT images. We found that the higher thickness of the membranes in the near-placental region is attributable to non-loadbearing layers (*i.e*., spongy layer, chorion, and decidua) and that the load-bearing amnion layer is not significantly different in thickness between the two anatomical regions (within the resolution and precision of our OCT imaging and segmentation). Therefore, the decrease in the strength of amnion in the near-cervical region must have a microstructural and biochemical source that is not reflected in measurements of its thickness.

**Fig. 6.**
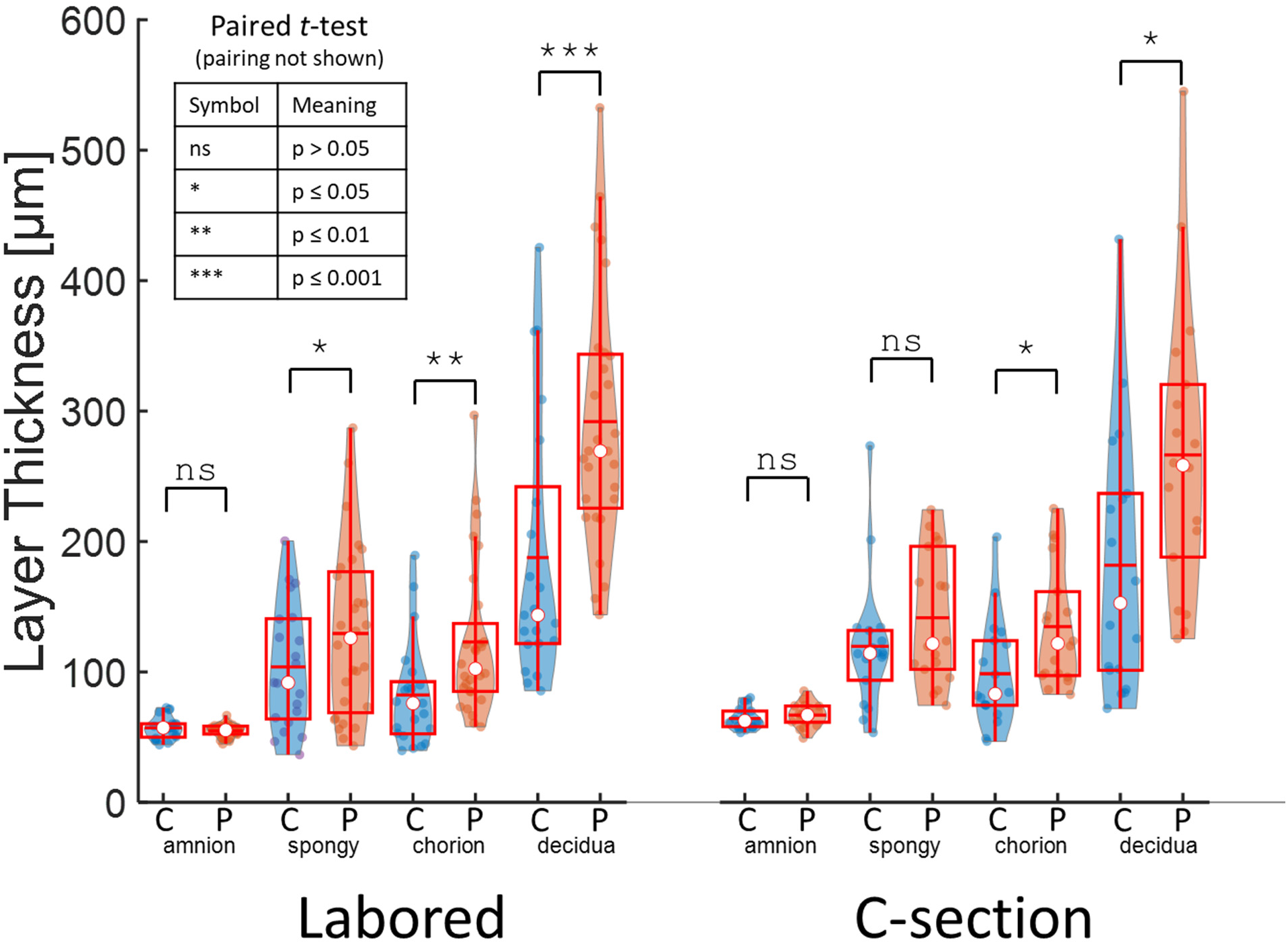
Non-load bearing layers account for location-dependent variability in thickness. Thickness of constituting layers of amniochorion, as estimated from the segmented OCT images, are compared between near-cervical (C) and near-placental (P) regions in both delivery groups. Amnion’s thickness, within the resolution and segmentation accuracy of OCT images, is not significantly different between the two locations. Other non-load bearing layers are significantly thicker in the near-placental region. Box plots show the interquartile range and whiskers extend from the box to 1.5 times the interquartile range. White dots show the median and red horizontal lines show the mean.

### 3.5 Tangent stiffnesses of fetal membranes near the cervical os are different from near the placental disk

Load response was investigated by analyzing the tension-strain loading curves of the samples from the two locations and comparing their tangent stiffness in the low-load toe region and the high-load linear region. Given the variable strain and tension ranges across samples, for a qualitative visual comparison of the loading curves from the near-cervical and near-placental locations, we normalized each loading curve strain and tension values by their respective maxima and plotted them in the same graph. Fig. 7 (a) and (b) show the normalized loading curves for the labored delivery group and the C-section group, respectively. In the labored delivery group (Fig. 7 (a)), samples from the near-cervical and near-placental locations group separately from each other. The near-cervical samples show a more linear loading curve, while the near-placental samples show a more non-linear elastic (*i.e*., more convex) loading curve. However, in the C-section delivery group (Fig. 7(b)), the near-cervical and near-placental samples do not show the same separation. There is also higher inter-sample variability in this group. For a more quantitative comparison of the sampling locations, tangent stiffness of the loading curves in the low-load toe region (Fig. 7 (c)) and the high-load linear region (Fig. 7 (d)) were plotted. In the labored delivery group, near-cervical samples compared to near-placental samples present with higher toe region tangent stiffness (*p*=0.006, paired *t*-test) and lower linear region tangent stiffness (*p*=0.045, paired *t*-test). However, these location-dependent trends do not hold for the C-section delivery group.

**Fig. 7.**
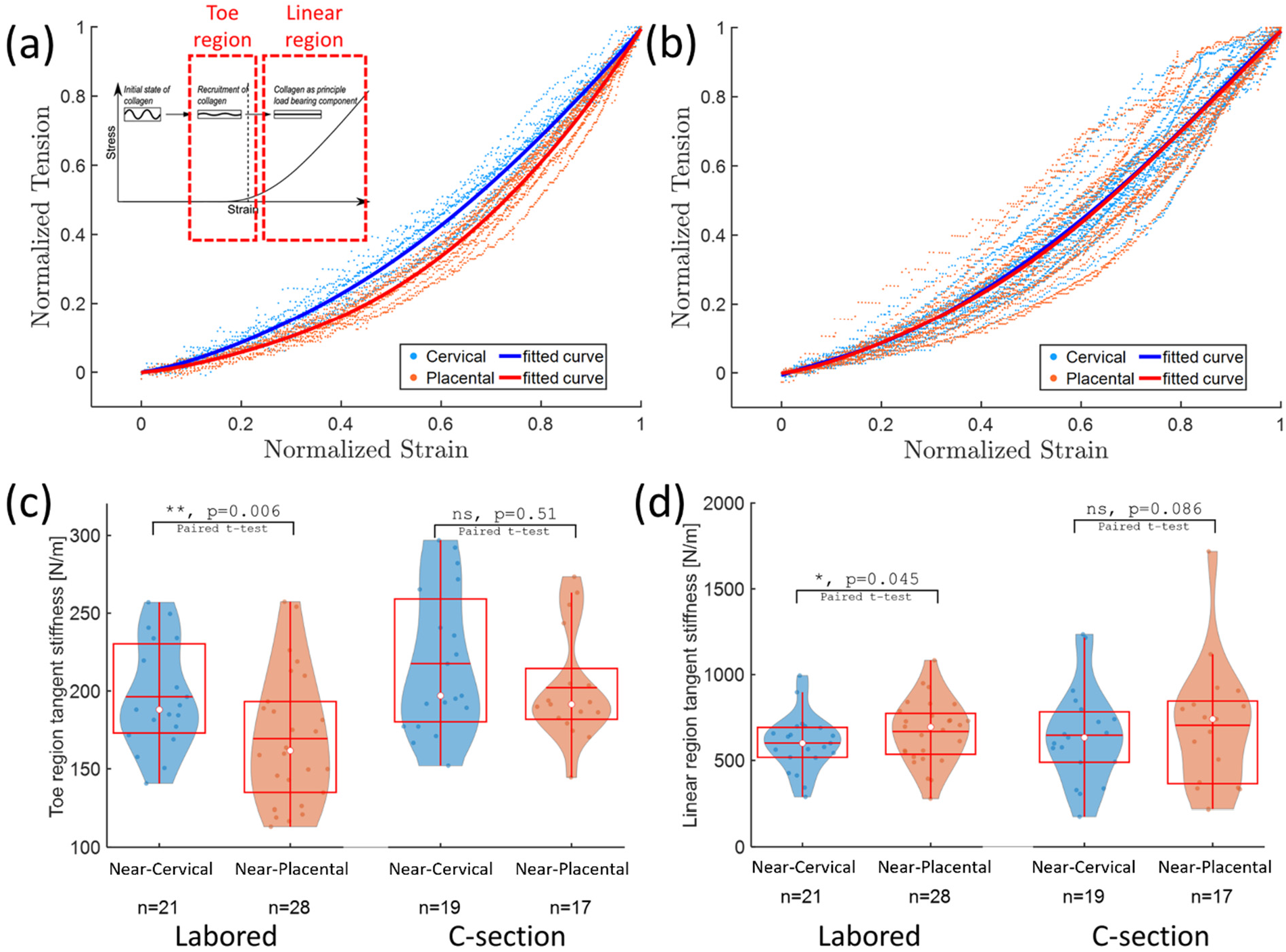
Cervical samples show signs of strain hardening and work fatigue. The loading curve of the composite amniochorion from near-cervical and near-placental locations are compared. The loading curve is divided into a toe region where collagen fibers are not fully tensioned, and a linear region where collagen fibers are tensioned and loaded until rupture. For a visual comparison, (a) shows the normalized loading curves of the labored delivery group samples and their fitted curves. The cervical samples show more linear loading curves that cluster separately from the placental samples with more nonlinear loading curves. The slope of the tension-strain loading curve (tension modulus) in either region is reported. The toe region was defined as the portion of the loading curve between 0.5 and 5 kPa of saline pressure, and the linear region was defined as the portion of the loading curve between 7.5 and 20 kPa of saline pressure. (b) shows the normalized loading curves and fitted curves for the C-section delivery group. Cervical and placental sample curves in this group do not cluster separately. (c) The tension modulus in the toe region is higher in the near-cervical location than the near-placental sampling location. However, the C-section group shows higher toe modulus than the labored group in both sampling locations and the location dependent difference does not reach significance in the C-section group. (d) The linear region modulus is higher in the near-placental location than the near-cervical location in the labored group, but the difference is less significant in the C-section group. Box plots show the interquartile range and whiskers extend from the box to 1.5 times the interquartile range. White dots show the median and red horizontal lines show the mean.

## 4 Discussion

This study presents the first application of OCT imaging for resolving the dynamic loading response of fetal membranes sub-layers from full-depth cross-sectional images along with application of a convolutional neural network for automated segmentation of the sub-layers. Previously, static OCT images of fetal membranes were compared to histology sections and features like obliterated chorionic villi, pseudo-cysts, and calcifications were identified, and thicknesses were measured [28–30]. Separately, mechanical studies of fetal membranes loading response to inflation by pressurized saline or aspiration by vacuum have evaluated mechanical properties of these tissues using video camera measurements of membrane deflections (either side view of apex rise or top view of ink markings) [24]. Here, we developed *ex vivo* methods to combine these approaches, specifically, to assess changes in sub-layer thickness within human fetal membranes during dynamic loading using an inflation test, volumetric OCT, and automated sub-layer segmentation with convolutional neural networks. Sources of contrast in OCT volumes were identified using corresponding histology and 2P microscopy and new neural networks were trained to identify layers within human fetal membranes under varying loading states. Differences in layer thickness and loading properties were compared between near-cervical and near-placental locations, and between labored and C-section deliveries.

Interestingly, unlike in the labored group, there was no significant difference in stiffness between the near-placental and near-cervical regions in the C-section group. This may be due to the manner in which the placenta and fetal membranes are manually extracted during C-section birth. Compared to spontaneous separation or controlled cord traction (CCT), manual removal may subject the placenta and fetal membranes to larger stresses particularly near the placental disk where adherence to the uterine wall is strongest [33]. As such, we hypothesize that these large stresses and the resulting plastic deformations may mask any pre-existing location-dependent mechanical differences between fetal membranes from near-placental and near-cervical regions in the C-section group. For example, rupture pressures of near-placental samples in the C-section group are smaller than the labored group (Fig. 5(b)) and are not significantly different from the near-cervical samples. The C-section near-placental amniochorion samples also show higher stiffness in the toe region of the loading curve (Fig. 7(c)) and are not significantly different from the near-cervical C-section samples, in contrast to the labored group amniochorion samples that show location-dependent differences in tangent stiffness. Therefore, it may be beneficial to control for the method of placenta extraction in future studies.

Prior studies indicated that overall thickness of the membranes is not correlated with their tensile strength [12], a finding replicated in our study (Fig. 5(d)). This confirms that, for an *in vivo* measurement of fetal membranes to provide diagnostically relevant information for PPROM risk assessment, resolution of the sub-layer breakdown of fetal membranes and their collagen matrix is necessary. Our results show that OCT provides the necessary resolution and contrast for such measurements. While our study was limited to term specimens due to difficulty of obtaining preterm samples in large numbers, the methods developed here may be applied in future studies to identify predictive markers of PPROM. The possibility of developing small-diameter fiber optic-based endoscopic OCT probes [34] for trans-cervical imaging of the fetal membranes further adds to the appeal of this technique for clinical use.

Overall, these studies fill a gap in our understanding of the structural and mechanical properties of human fetal membranes at high resolution under dynamic loading events, and inspire further study, particularly *in vivo*.

## Supporting information

Visualization 1

Visualization 2

## Acknowledgements

We acknowledge funding from the Morgridge Institute for Research (KS and MCS), Carol Skornicka Chair of Biomedical Imaging (MCS), Retina Research Foundation (RRF) Daniel M. Albert Chair (MCS). We thank the Birthing Center at the UnityPoint Heath Meriter Hospital and the Intermountain Healthcare Utah Valley Hospital Labor and Delivery for their support in sample collection. Illustration cartoon was created with BioRender.com.

## Disclosures

The authors declare no conflicts of interest.

## Data availability

Data underlying the results presented in this paper are not publicly available at this time but may be obtained from the authors upon reasonable request.

## Notes

### Competing Interest Statement

The authors have declared no competing interest.

